# The collective application of shorebird tracking data to conservation

**DOI:** 10.1101/2024.01.30.576574

**Authors:** Autumn-Lynn Harrison, Candace Stenzel, Alexandra Anderson, Jessica Howell, Richard B. Lanctot, Marley Aikens, Joaquín Aldabe, Liam A. Berigan, Joël Bêty, Erik Blomberg, Juliana Bosi de Almeida, Andy J. Boyce, David W. Bradley, Stephen Brown, Jay Carlisle, Edward Cheskey, Katherine Christie, Sylvain Christin, Rob Clay, Ashley Dayer, Jill L. Deppe, Willow English, Scott A. Flemming, Olivier Gilg, Christine Gilroy, Susan Heath, Jason M. Hill, J. Mark Hipfner, James A. Johnson, Luanne Johnson, Bart Kempenaers, Paul Knaga, Eunbi Kwon, Benjamin J. Lagassé, Jean-François Lamarre, Christopher Latty, Don-Jean Léandri-Breton, Nicolas Lecomte, Pam Loring, Rebecca McGuire, Scott Moorhead, Juan G. Navedo, David Newstead, Erica Nol, Alina Olalla-Kerstupp, Bridget Olson, Elizabeth Olson, Julie Paquet, Allison K. Pierce, Jennie Rausch, Kevin Regan, Matt Reiter, Amber M. Roth, Mike Russell, Sarah T. Saalfeld, Amy L. Scarpignato, Shiloh Schulte, Nathan R. Senner, Joseph A. M. Smith, Paul A. Smith, Zach Spector, Kelly Srigley Werner, Michelle L. Stantial, Audrey R. Taylor, Mihai Valcu, Walter Wehtje, Brad Winn, Michael B. Wunder

## Abstract

Addressing urgent conservation issues, like the drastic declines of North American migratory birds, requires creative, evidence-based, efficient, and collaborative approaches. Over 50% of monitored North American shorebird populations have lost over 50% of their abundance since 1980. To address these declines, we developed a partnership of scientists and practitioners called the Shorebird Science and Conservation Collective (hereinafter “the Collective”). Here, we present this successful case study as an example for others engaged in translational science. The Collective acts as an intermediary whereby dedicated staff collate and analyze data contributions from scientists to support knowledge requests from conservation practitioners. Data contributions from 74 organizations include over 6.7 million shorebird locations forming movement paths of 3,345 individuals representing 36 species tracked across the Americas. We describe the founding and structure of the Collective and conservation activities we supported in our first two years. As the volume of scientific data on animal movements continues to grow, groups like the Collective can be vital liaisons to rapidly integrate and interpret research to support conservation action.

## Introduction

Conservation efforts are often location-specific (Stewart et al. 2013), presenting challenges for managing migratory species that traverse vast distances. Shorebirds (Charadriiformes, suborder Charadrii) exemplify these challenges as many travel thousands of kilometers biannually across continents and oceans (e.g. Gill et al. 2009). The flocking behavior of shorebirds gives the illusion of abundance, leading to shifting baselines (*sensu* Pauly 1995) and masking major declines. Shorebirds have the largest declines (net and % loss in abundance; greatest proportion of species in decline) of nearly 3 billion birds lost in North America since 1970 (compared to landbirds, waterbirds, and waterfowl, Rosenberg et al. 2019). Declines show no sign of abating (Smith et al. 2020), emphasizing the urgent need for evidence-based conservation action for shorebirds. Here we present a partnership of scientists and practitioners (defined here as those directly guiding conservation activites) responding to this need for action by linking a relatively new scientific resource—shorebird tracking data—with conservation practice.

Scientists worldwide use electronic tracking devices (hereinafter “tags”, reviewed by Robinson et al. 2010) to collect animal movement data (hereinafter “tracking data”). Tracking data have become instrumental in unveiling migration routes; identifying important habitats; inferring animal behavior, and understanding population connectivity (Kays et al. 2015; Hussey et al. 2015). Aggregating tracking data across species has led to significant insights, for example documenting patterns in timing of seasonal events in relation to climate change in the Arctic (Davidson et al. 2020). Insights from tracking data have also led to significant conservation achievements. For example, tracking data collected from marine animals were used to designate marine protected areas, reduce fisheries bycatch and vessel strikes, and minimize impacts of extractive industries (Hays et al. 2019).

There has been significant growth in shorebird tracking studies over the last 20 years, unlocking the potential for tracking data to contribute to shorebird conservation. In a review of avian tracking studies (Scarpignato et al. 2023), shorebirds and songbirds constituted <1% of studies before 2006, but these groups constituted 30–75% of tracking studies in subsequent years. However, the conservation potential of these increasingly abundant data is limited by four major analytical and conservation challenges (Table 1): only 35% of shorebird tracking data are open access (Scarpignato et al. 2023); analyzing tracking data requires specialized knowledge; shorebirds are not the management priority in many of their important habitats; and due to their migratory nature, there is a need for strategic integration across organizations and habitats.

**Table 1.** Factors limiting the application of shorebird tracking data to conservation include limitations to science accessibility and analysis, and limitations to conservation implementation.

|  |  |  |
| --- | --- | --- |
| Science Limitations | **Data Access**<br>Finding and securing access to relevant data is time-consuming<br>• Despite increasing use of data repositories (e.g. Movebank, Kays et al. 2022), only 35% of shorebird tracking datasets (reviewed by Scarpignato et al. 2023) were accessible for conservation.<br>• Within repositories, data are spread across hundreds of unique studies led by many scientists.<br>• Obtaining private data involves asking permission from many owners. | **Data Analysis**<br>Analyzing tracking data for applications is not straight-forward<br>• Analyzing tracking data requires specialized statistical and software knowledge (O'Toole et al. 2021; Joo et al. 2020; Gupte et al. 2022).<br>• Data integration challenges are common in ecology and limit the use of datasets for the public good (Reichman et al. 2011).<br>• Practitioners typically do not have analysis as a primary job function (Blickley et al. 2013). |
| Implementation Limitations | **Competing Priorities**<br>Shorebirds are not the priority within many of their important habitats<br>• Although interior wetland management in North America has moved from a waterfowl focus to an “all birds” approach, the extent to which shorebirds are integrated into wetland planning still varies (Doherty et al. 2018).<br>• Many shorebirds rely on working lands where management may focus on food commodities production or on game or economically important species.<br>• Some management programs need updating to reflect changing conservation circumstances (Sutherland et al. 2012).<br>• Practitioners come from diverse professional backgrounds (Fernández 2016; Ciuzio et al. 2013) that may not include a background in shorebird ecology, or how to manage for them alongside primary objectives. | **Need for Strategic Integration**<br>Diverse groups undertake shorebird conservation<br>• Groups work at all geographic scales (Table 3), have varied funding mechanisms, operate under a variety of legal frameworks, and use varied approaches to science-based conservation implementation.<br>• Within groups, distinct individuals can be responsible for research, regulations, program development, or technical assistance.<br>• The lack of a central scientific resource focused on shorebirds limits the ability to integrate knowledge across groups to help target local actions to support hemispheric priorities (Brown et al. 2001). |

To overcome challenges limiting science translation for conservation, both linear and interactive approaches are used (Beier et al. 2017; Toomey et al. 2017; Saunders et al. 2021). Linear methods involve creating scientific products to be accessed by practitioners (e.g. data portals, online tools, publications, and manuals). Despite conservation benefits (Seavy & Howell 2010; Sullivan et al. 2017), linear delivery has limitations. Tools may not suit practical purposes due to formatting or language incompatibilities, incomplete data, or mismatched questions, and can be time-consuming to find and synthesize (Vogel et al. 2007). In contrast, interactive approaches range from focused workshops to long-term co-production where research and products are collaboratively designed, planned, and implemented by practitioners and scientists (Young et al. 2014; Roux et al. 2006; Saunders et al. 2021). These approaches allow participants to ask clarifying questions (Gerber et al. 2020), build and maintain relationships (Lindell & Dayer 2022), and engage in active collaborations (Young et al. 2014). Interactive approaches are generally more successful (Cooke et al. 2020), but there are sometimes participation barriers (Seavy & Howell 2010; Oliver et al. 2019).

We combined elements of both linear and interactive knowledge-sharing approaches to form the Shorebird Science and Conservation Collective (hereinafter, ‘the Collective’). The Collective is a partnership of scientists and practitioners working to advance shorebird conservation in the Americas (Figure 1). Our goal in sharing the origin, structure, and achievements of the Collective is to provide a case study that can help guide others who are interested in increasing the impact of existing science on conservation. The Collective acts as an intermediary (or boundary group, Cash et al. 2006) whereby dedicated staff collate and analyze data contributions from scientists to support the conservation needs of practitioners. We describe how the Collective a) raised funds, b) initiated a focused proof-of-concept, c) requested data, established processes to address privacy concerns and to report to contributors, d) developed a co-production process to listen and respond to the needs of practitioners, and e) contributed to conservation in its first two years.

**Figure 1.**
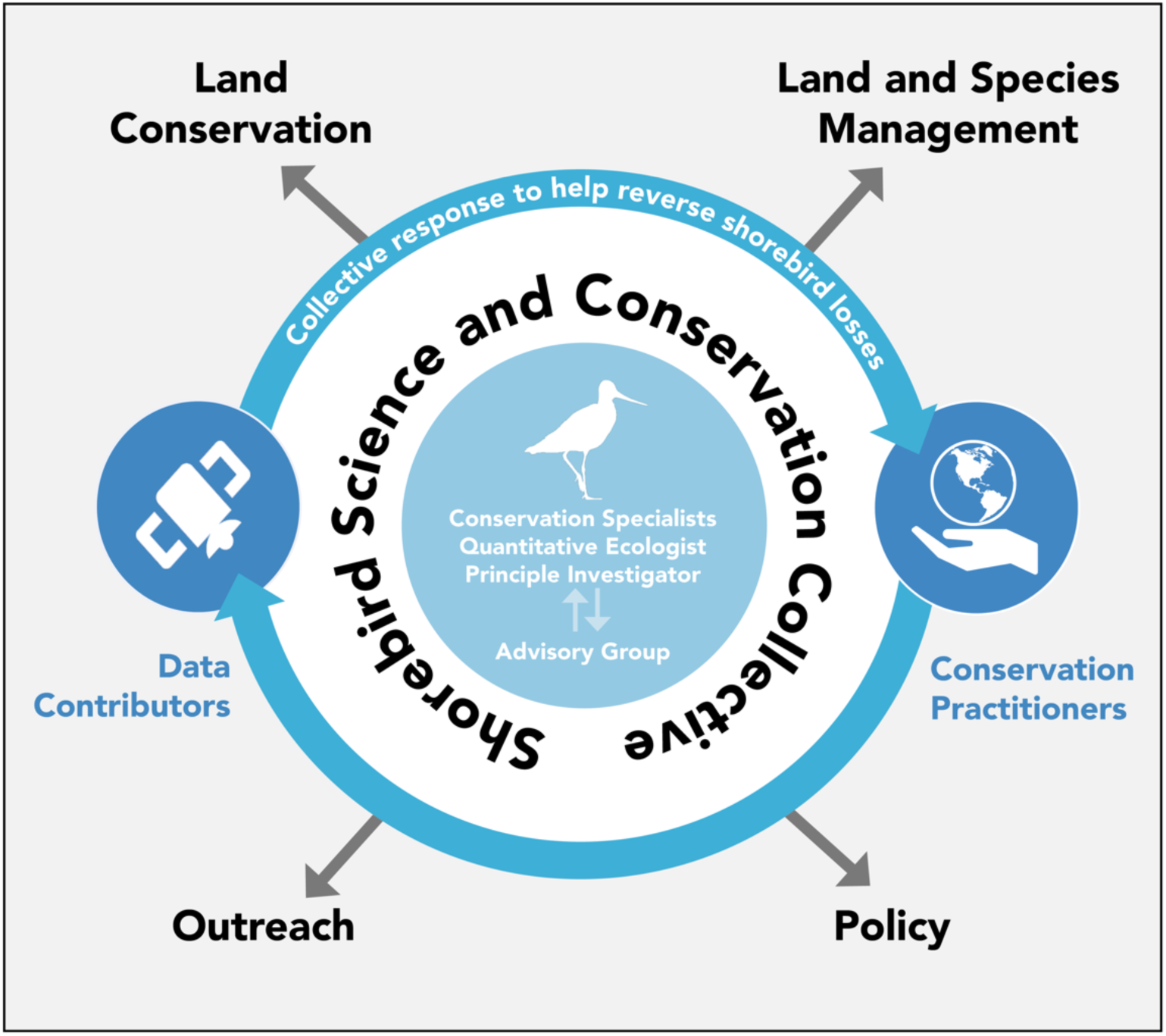
The process by which the Shorebird Science and Conservation Collective distributes knowledge generated from electronic tracking studies of shorebirds to conservation partners across the Americas to enhance land and species conservation, management, and outreach and education activities. In the future, this resource also has potential to inform conservation policy. The Collective engages with practitioners in service to community-driven management and conservation needs, with the ultimate goal of reversing population declines.

### Forming the idea for the Shorebird Science and Conservation Collective

Private foundations can greatly impact conservation funding (Zavaleta et al. 2008; Bakker et al. 2010). In our case, the Knobloch Family Foundation catalyzed the Collective by inviting the Principal Investigator to propose a “transformative idea for data-driven shorebird conservation in North America.” The idea to collate shorebird tracking studies for conservation was inspired by the successful example of BirdLife International’s Seabird Tracking Database in informing conservation (BirdLife International 2004)—but adapted for unique considerations of how and where shorebird conservation is implemented (Table 1, Table 3). Given existing limitations (Table 1), a linear approach facilitating data discovery or access, but requiring practitioners to interpret data and literature themselves, seemed unlikely on its own to maximize knowledge transfer. Instead, we proposed a Collective that would extend beyond a repository to include staff for data analysis and co-development of conservation products to support direct requests from practitioners.

A funding proposal was developed with feedback from shorebird researchers and conservation practitioners through open-invitation webinar discussions. Many participants were already sharing, co-developing or applying research to conservation. As conceived, the Collective would add value by expanding conservation opportunities, rather than compete with or replace existing partnerships. Webinars were instrumental for assessing support, refining ideas, and identifying existing gaps in translating science to shorebird conservation.

### A proof-of-concept for the Collective

We proposed a three-year proof-of-concept focused on the midcontinent of North America to evaluate the Collective’s structure and gauge demand for shorebird tracking data. The Midcontinent has been elevated as a priority landscape by multiple conservation entities (www.grasslandsroadmap.org, www.midamericasshorebirds.org) and provides habitat for 42 of the 50 shorebird species that regularly breed in North America (Donaldson et al. 2000; Brown et al. 2001). For example, the Texas Gulf Coast provides migratory and wintering habitat for more than 1 million shorebirds annually (Withers 2002) while the Prairie Pothole Region’s wetlands and grasslands (Doherty et al. 2018) support millions of migrants each spring and fall (Skagen 2006; Steen et al. 2018). These landscapes have experienced major wetland loss (up to 90% at the U.S. state level, (Dahl 1990; Dahl 2014)) and have high spatial and temporal climatic variability from droughts to flooding, making cohesive shorebird management challenging across the region and among years (Fellows et al. 2001; Russell et al. 2016; Skagen & Thompson 2013). Many areas are privately owned (e.g. nearly 90% of the Great Plains is non-public (NRCS 2021), requiring coordination with landowner-focused agencies, which deliver voluntary incentive programs and/or provide technical assistance to manage habitat on working lands (e.g., NRCS, USFWS Partners for Fish and Wildlife Program), as well as state and local governments, land trusts, and other conservation entities.

The proposal was funded in late 2020, allowing implementation to begin in early 2021.

### The Collective’s structure and systems

The Collective is comprised of 4 groups of people (Figure 1): a core operational team (P.I., staff, and Chair of the Advisory Group), an Advisory Group (Appendix I), data contributors, and practitioner partners. All groups are represented as co-authors. The P.I. and staff are based at the Smithsonian Migratory Bird Center. Staff include a quantitative ecologist who aggregates, processes, and analyzes data; and conservation specialists who work with practitioners to inform them of the tracking data resource, assess their information needs, and help prepare conservation products. The 16-person Advisory Group advises on maximizing the conservation benefit of the Collective, promotes its value to potential data contributors, and connects it with organizations engaged in similar efforts or those that would benefit from tracking data. Data contributors typically are researchers from within public, private, or non-profit organizations. Practitioners directly guide conservation actions from within management agencies, trusts, and non-profit groups (Table 3). Some data contributors are also practitioners.

With guidance from all involved, the core team developed clear operational processes including a Data Sharing Agreement (Appendix II), a conservation use request form (Appendix III), and a conservation reporting system (Appendices IV-VI) to ensure data privacy agreements are upheld and to increase accountability to data contributors and practitioners.

#### Crafting a Data Sharing Agreement

Data contributors are the Collective’s foundation; their data drive evidence-based recommendations and products. Feedback indicated that careful crafting of a Data Sharing Agreement would ensure maximum trust and participation by the scientific community. The Collective aims for shorebird tracking data to be findable, accessible, interoperable, and reusable (FAIR Guiding Principles, Wilkinson et al. 2016) while respecting contributor rights. Many factors influence data contribution (e.g., Roche et al. 2020), with some policies mandating public sharing. Conversely, certain datasets, like recently-collected data or those restricted by endangered species status, may require embargo or organizational approval before public release. We thus drafted a flexible agreement (Appendix II, https://nationalzoo.si.edu/migratory-birds/shorebird-science-and-conservation-collective-data-sharing-agreement-form). All contributors allow the Collective’s core team to explore data to assess applicability to a conservation request. Contributors further customize usage permissions (either providing full permission or requiring a permission request before each use) within use categories: conservation, education, funder reports, scientific publications, and/or research requests.

#### Compiling data

We invited data contributions through email, a webinar, social networks, and word-of-mouth. Data were received either through repositories (e.g., Movebank) or were sent directly to the Collective’s staff. We prioritized processing data with higher spatial accuracy (e.g., Argos and GPS; Table 2) that we anticipated would be most relevant to conservation requests from practitioners. We developed a set of R scripts that first apply pre-filters (e.g. to determine when a tag has become stationary, indicating it has either been shed by the bird or the bird has died).

**Table 2.** Description and sample size of data types submitted to the Shorebird Science and Conservation Collective as of October 2023.

| TECHNOLOGY | DESCRIPTION | DATA FORMAT | SPATIAL ACCURACY | SAMPLING FREQUENCY | MONITORING DURATION | NUMBER OF INDIVIDUALS CONTRIBUTED (% of individuals) | NUMBER OF SPECIES CONTRIBUTED (% of species) |
| --- | --- | --- | --- | --- | --- | --- | --- |
| **GPS** | Electronic tags attached to individual birds. Locations estimated from GPS satellites and obtained either by retrieving the device and downloading stored data (i.e., archival tags) or transmitting the data to satellites. | Latitude and longitude coordinates | Typically 10-100 m (Noonan et al. 2019) | Minutes to days or pre-programmed schedules such as every 2 weeks during overwintering and 2 days during migration depending on study goal and tag battery life | Months to years depending on tag type and attachment method | 1,106* (33.1 %) | 17 (47.2 %) |
| **Argos Doppler Shift** | Electronic satellite tags (called Platform Terminal Transmitters or PTTs) attached to individual birds. Tags send signals to orbiting satellites that re-transmit location data back to a receiving station on Earth for processing. | Latitude and longitude coordinates with “location class” codes or error ellipses that indicate the accuracy of the estimated location | Generally 250—1,500 m (CLS America 2016) but individual locations can have errors larger than 25km (Jonsen et al. 2020) | Minutes to days, often irregular; tags may use pre-programmed on/off schedules to allow for charging (solar models), or to conserve energy (battery models) | 1+ years depending on tag type and attachment method | 853* (25.5 %) | 14 (38.9%) |
| **Motus** | Electronic VHF radio tags attached to individual birds that | Raw detection data (e.g., timestamp, | Typically 15 – 20 km from receiving stations (Taylor et | 3 - 30 seconds when tag nears receiving station, | Days to months | 1,091 (32.6 %) | 4 (11.1%) |
|  | are detected by a network of receiving stations. Detection data are stored on receivers, synched over the internet, or sent via satellite or cellular network. | signal strength) associated with the location of each receiving station | al. 2017) but up to 180 km (Anderson et al. 2019); ~500 m if omnidirectional antennas are used (Taylor et al. 2017) | generally, though gaps can be minutes to days |  |  |  |
| **Light-level Geolocator** | Electronic devices attached to individual birds that record ambient light levels, which can be compared to global light patterns to estimate the animal's location. Devices must be retrieved to download the data. | Raw ambient light levels which require substantial processing and analysis to estimate locations (Lisovski et al. 2020) | 50 - 200 km with much greater error in latitude during spring and fall equinox (Lisovski et al. 2012; Rakhimberdiev et al. 2016) | 1-2 locations per day | 1+ years | 483 (14.4 %) | 15 (41.7 %) |
| **Resight** | Observations of individual birds marked with unique color band / flag combinations, or alphanumeric bands or flags | Latitude and longitude coordinates provided by the observer | Depends on observer proximity but usually < 100 m. | Limited to where and when observers search for birds, marker retention, and animal lifespan | Limited to observer effort, marker retention, and animal lifespan | Submitted data currently not integrated into the Collective dataset | 15 (41.7%) |
|  |  |  |  |  |  | * Includes 188 individuals tracked with both GPS and Argos technologies |  |

**Table 3.**
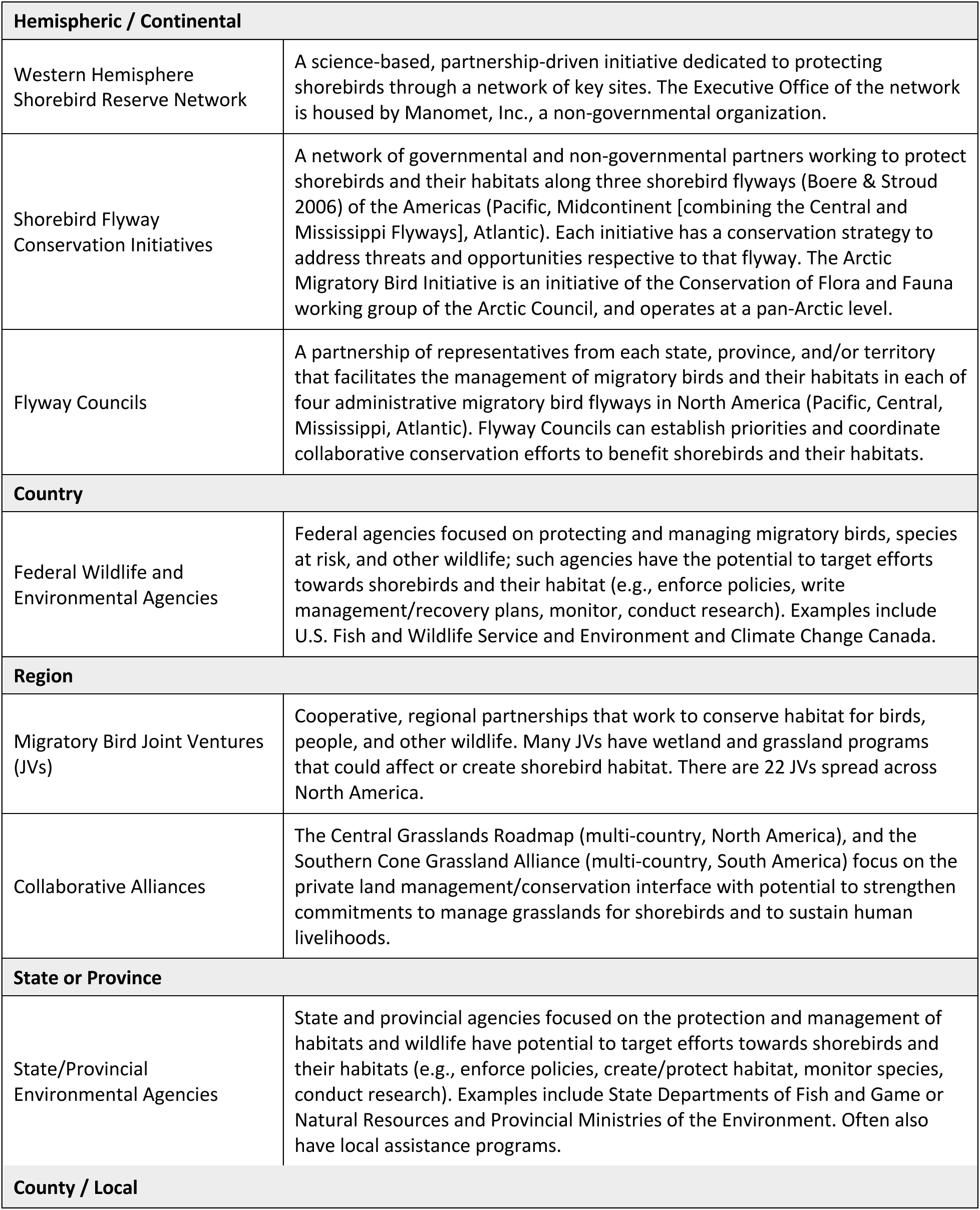
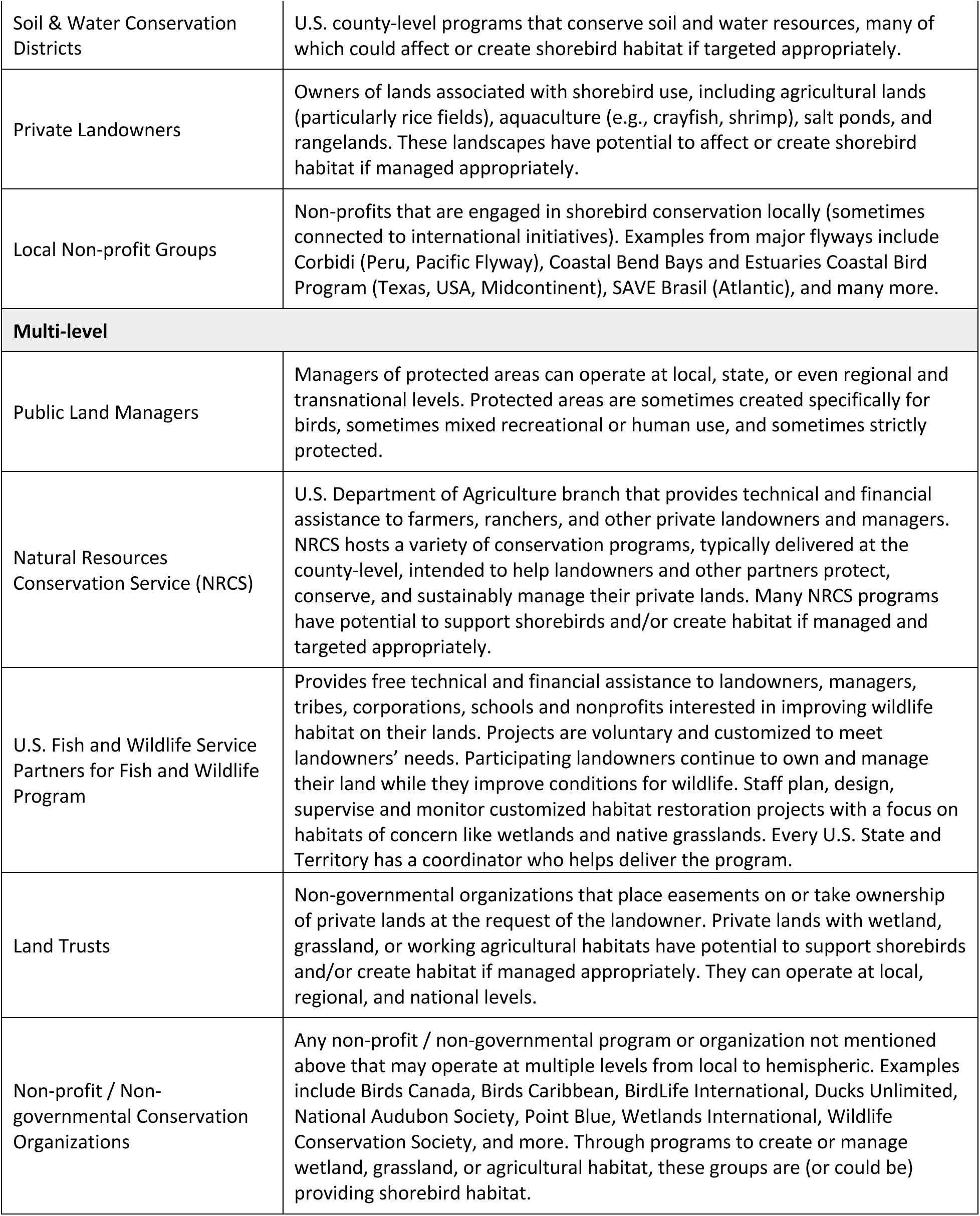
Examples of organizations, initiatives, and individuals engaged in shorebird conservation and management and the geographic scale (hemispheric to local) at which these organizations operate.

Then a continuous-time state-space model (Jonsen et al. 2023) estimates the most probable movement path accounting for the 1) spatial error of the tracking technology and 2) irregular sampling across individuals and species. The Collective is currently integrating data collected by the Motus Wildlife Tracking System (Taylor et al. 2017) and light-level geolocators (Table 2) that were contributed either pre-processed using various methods, or in raw, unprocessed form, requiring additional processing or standardization to ensure appropriate interpretation (Lisovski et al. 2018). Data are linked to their associated Data Sharing Agreements to ensure that permissions and attributions are closely followed by staff when considering use for conservation applications.

#### Reporting conservation uses to contributors

Contributors are notified when their data are relevant to a conservation request (examples in Appendices III-V). The notification describes the project and provides a map of the data involved. For any data use that the contributor did not pre-approve in the Data Sharing Agreement, the Collective requests permission before proceeding. While this process adds to the administrative work of the Collective’s staff, it incentivizes and rewards data contribution, increases accountability, safeguards privacy settings, and promotes dialogue to prevent misinterpretations of contributed data.

#### Engaging conservation partners

Information requests from practitioners are submitted via a Conservation Use Request Form (Appendix III and https://nationalzoo.si.edu/migratory-birds/request-for-use-shorebird-tracking-data), or through emails, informational meetings, conference presentations, and referrals. Conservation specialists currently prioritize outreach to practitioners within the Midcontinent given the geographic focus of our proof-of-concept. However, requests have been received from across North America. Knowledge-sharing typically begins with a 1-hr virtual meeting between practitioners and members of the Collective’s core team. The team describes the Collective’s approach for data-driven conservation, and practitioners describe their work, priorities, and shorebird-related questions. If there is opportunity for the Collective to inform a project, further conversations help refine questions, information needs, and preferred formats of data products. Data contributors are then engaged to notify them that their data are relevant to a conservation request and/or to request further permissions.

Throughout the project, the core team communicates with practitioners and data contributors as needed. The team presents findings to practitioners, allowing questions and feedback regarding the delivery format and recommendations. At the project’s conclusion, a Conservation Contribution Report summarizes recommendations, acknowledges data contributors, and provides information about shorebird ecology and best management practices. A public version (with any sensitive data removed) is also shared online (https://nationalzoo.si.edu/migratory-birds/shorebird-collective). The Collective classifies conservation contributions according to the IUCN Conservation Actions Classification Scheme (Salafsky et al. 2008; IUCN 2012) to align with standard conservation evidence reporting (Sutherland et al. 2019).

### The wealth of shorebird tracking data available for conservation in the Americas

Data contributions include 6.7 million locations from 3,345 individual shorebirds of 36 species (Figure 2). Since January 2021, when data were first requested, contributors have signed 58 agreements for data originating from 74 organizations (e.g., academic institutions, nonprofits, and government agencies) and 136 data co-owners from 14 countries.

**Figure 2.**
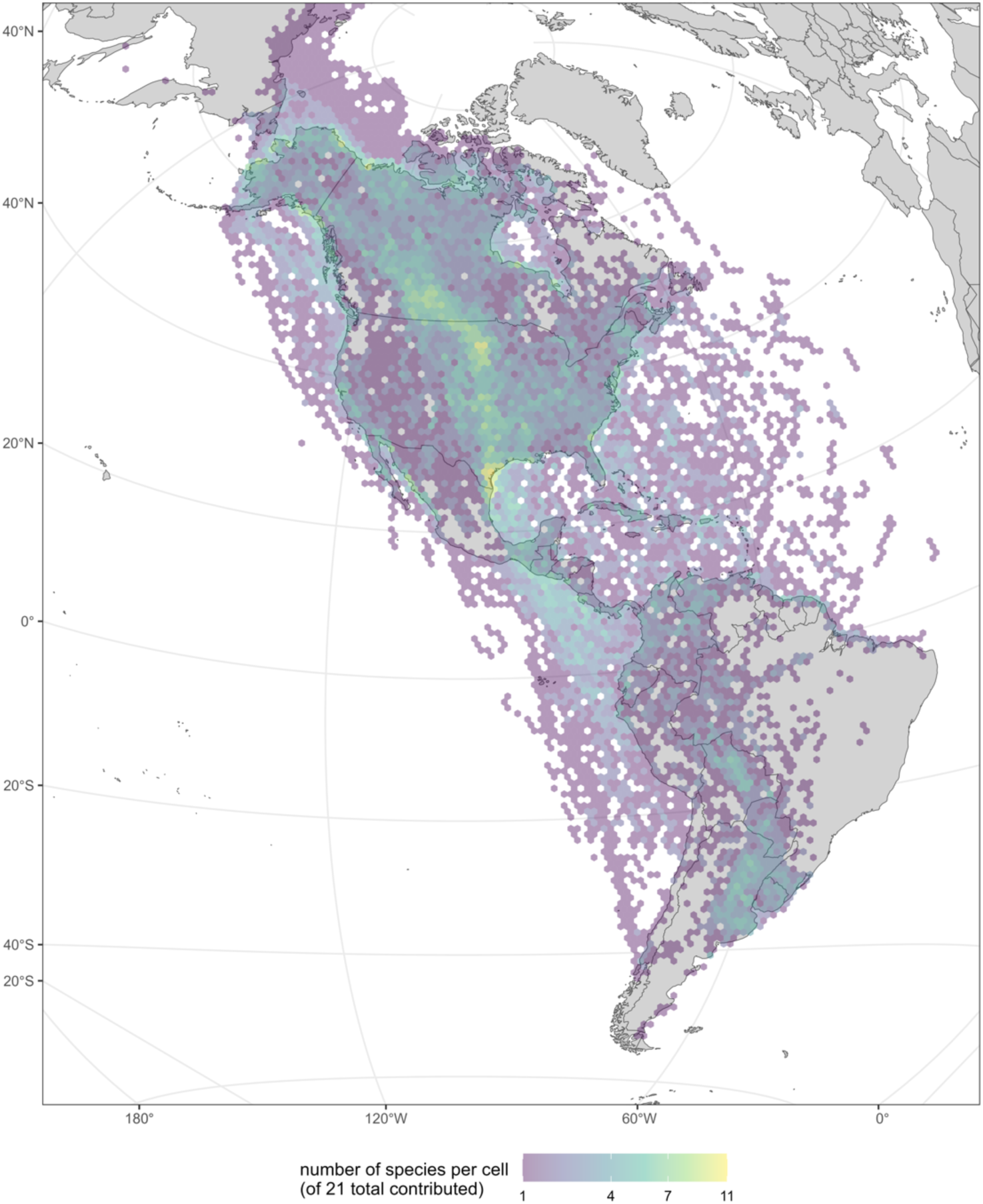
Distribution of Argos and GPS tracking data (Table 2; n=1,771 individuals) contributed to the Shorebird Science and Conservation Collective as of October 2023, shaded by the number of species with tracking locations in 100 km x100 km hexagonal cells.

Data span the Americas with areas in the Midcontinent demonstrating clear importance to multiple species, including the Prairie Pothole Region and Gulf of Mexico coast (Figure 2). Shorebirds were tracked via light-level geolocator, Argos, GPS, and Motus technologies, as well as with visual observations of uniquely marked birds (i.e. with flags or bands) (Table 2, Figure 2). Data come from the smallest (Least Sandpiper, *Calidris minutilla,* 19-30g, tracked with 0.67 g Motus tags; (Anderson et al. 2019)) to the largest (Long-billed Curlew, *Numenius americanus,* 490-950 g, tracked with 9.5-22 g satellite tags; (Page et al. 2014)) shorebirds in North America (Species list, Appendix VII). Detections came from varied habitats including fresh and saline wetlands, uplands, grasslands, agricultural fields, coastal beaches and mudflats, tundra, forest, and over open ocean.

### Contributing to conservation

Since our first conservation request in November, 2021 to October, 2023, the Collective has supported 10 conservation projects (with another 4 ongoing). We have sent 93 notifications to 23 contributors informing them that their data collected from 17 species were relevant to conservation requests. Fifty-six notifications were for datasets pre-approved to be used for conservation, and 37 were requests to contributors requiring further permission (see Appendices II-IV for examples). Of the latter set, 92% were approved. Below, we provide examples of ways the Collective applied tracking data to support: 1) land conservation, 2) land and species management, and 3) education and outreach efforts.

## 1. Private Land Conservation

Both public and private lands support shorebird conservation in North America, but within the Midcontinent where there is high private ownership, private lands are an especially crucial component of environmental stewardship and land protection (Merenlender et al. 2004).

Trusts, foundations, and government agencies can use insights from tracking data to help guide conservation investments and secure funds for voluntary land purchase or conservation. These datasets are especially useful when they provide the only evidence of shorebirds using areas (for example, when systematic ground surveys or eBird observations are unavailable).

### Funding and siting of conservation easements

The Texas Parks and Wildlife Foundation’s Buffer Lands Incentive Program (BLIP) provides financial assistance for conservation easements (Merenlender et al. 2004) on privately owned “buffer lands” bordering Texas Parks and Wildlife Department (TPWD) public lands.

The BLIP requested evidence for shorebird use of buffer lands bordering 12 priority TPWD lands as one metric in their rubric to prioritize applications for conservation easement funding; half of BLIP funds are allocated for shorebird and grassland bird projects. Because many parcels under consideration are inaccessible to the public, direct evidence of use by shorebirds comes almost exclusively from tracking data. Fifty-two individuals of nine species in the Collective’s dataset stopped or overwintered on buffer lands. For 83% of individuals, tags were deployed out-of-state. The Collective summarized when birds stopped or wintered in buffer lands, the habitats used, and global migration links ranging from Russia to Argentina to assist with landowner engagement. Finally, we shared a list of all TPWD lands visited by tracked shorebirds in the Collective’s dataset, and state-wide density maps of individuals and species tracked.

### Purchasing land for conservation

The Conservation Fund (TCF) and The Nature Conservancy (TNC), two non-governmental conservation organizations, separately sought information to support funding efforts to acquire lands for permanent protection in the Texas coast (TCF) and the Texas panhandle (TNC). Although no shorebirds within the Collective’s dataset visited the parcels under consideration, 23 individuals of four species were tracked nearby (within 3–18 km). We used habitat layers and standardized eBird relative abundance (Fink et al. 2022) to provide additional context about the potential value of the parcels to shorebirds. Some habitats in the parcels seemed promising and we recommended on-the-ground surveys to confirm shorebird use. Both organizations shared these data with funders to demonstrate the potential value of these landscapes for shorebirds.

## 2. Land and Species Management

Many shorebird conservation activities manage for habitat or species of concern (Iglecia & Winn 2021). Land management includes activities like deploying water to create habitat, alternating wetting and drying of rice fields timed with shorebird usage, and prescribed grazing and burning of grasslands or rangelands. Species management includes activities like species recovery planning and status assessments, harvest management, and beach closures. Tracking data can reveal important sites, connectivity among sites, individual arrival and departure timing at particular sites, and species population structure to inform management actions.

### Informing the timing and location of freshwater deployments

The Galveston Bay Foundation and Texas Water Trade, two Texas-based non-governmental organizations, recently purchased over 5,000 acre-feet of water rights. They wanted to know when and where to deploy water along the east Texas coast to benefit shorebirds. “Focused flows” offer many benefits including habitat creation and restoration, salinity management, and continued crop yields during drought (Culp et al. 2014; Garmany 2020). Tracking data from six shorebird species provided information on local habitat use and timing through the focal area (Figure 3a, 3b). Standardized relative abundance data from eBird (Fink et al. 2022) added insights for species with limited tracking data and provided peaks of occurrence across multiple species (Figure 3b). We provided partners with recommendations for when to deploy water (preferred and alternate times, Figure 3c) and shorebird habitat management best practices.

**Figure 3.**
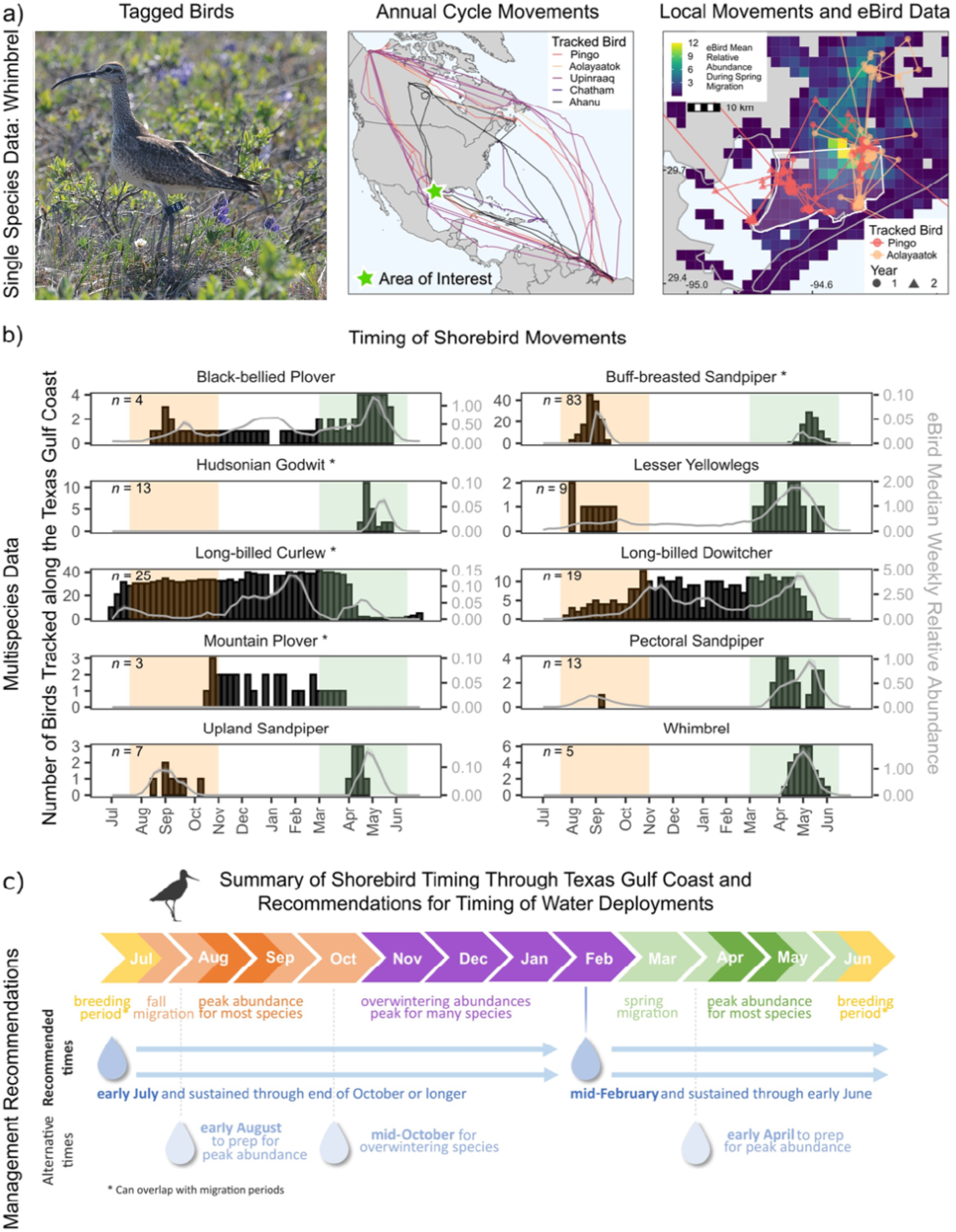
Example figure from a conservation project report informing the timing of freshwater deployments to provide habitat for migratory and wintering shorebirds in Texas. A) Example tracking data from Whimbrel, shared by Jennie Rausch, Environment and Climate Change Canada (middle) and overlaid on a focal area of interest in Texas (right). B) Time series of when tracked birds traveled through the Texas Gulf Coast (black histogram bars) together with median weekly relative abundance (gray lines) in the area of interest predicted by eBird community science data (Fink et al. 2022). Individuals tracked in more than one year contributed to histogram counts more than once because timing varied for individuals between years. Fall migration is shaded orange and spring migration is shaded green. Species with asterisks are Texas State Species of Greatest Conservation Need (Texas Parks and Wildlife Department 2012). Information from A) and B) were used to determine time periods during which water deployments could benefit peak diversity and abundances of migratory shorebirds, leading to C) Recommendation to conservation partners for deploying water to benefit migrating and overwintering shorebirds under recommended and alternate scenarios.

### Offshore wind energy development

Wind energy is important for reducing carbon emissions, but may have cumulative negative effects on birds through collisions or displacement (Goodale & Milman 2016; Fox & Petersen 2019). Tracking data can provide valuable insight about movement routes and timing of passage in proposed and active offshore wind areas to better understand shorebird vulnerability, inform siting decisions, and/or encourage “smart curtailment” strategies during high-risk migratory periods (Loring et al. 2021; Schwemmer et al. 2023). The Canadian Wildlife Service, Canada’s federal wildlife agency, requested information for two rapid (18-month) regional assessments offshore Nova Scotia and Newfoundland and Labrador to understand the temporal and spatial distribution of shorebirds moving through the focal areas. We provided maps of 41 individuals of eight species that migrated through the offshore areas of interest and highlighted areas of high activity (i.e., where track lines congregated for multiple individuals/species). The U.S. Fish and Wildlife Service, the United States’ federal wildlife agency, has requested similar data to incorporate into a collision risk model and to inform official comments on environmental impact statements. The Collective is actively supporting this request.

### Improving length of stay estimates and bird use days

Population abundance and trends are critical metrics that guide conservation actions and targets. However, accurate estimates are challenging to obtain for many shorebird species that aggregate in large groups with regular inflow and outflow of individuals (Vallecillo et al. 2021; Farmer & Durbian 2006; Gillings et al. 2009). Tracking data provide reliable estimates of length of stay (time between arrival and departure) for individuals at migratory stopover sites that can improve survey-based estimates of passage populations (Farmer & Durbian 2006). This information in turn, facilitates population monitoring, guides habitat management objectives and helps assess management effectiveness, and informs other planning tools such as abundance targets (Environment and Climate Change Canada 2016; U.S. Fish and Wildlife Service 2023) and harvest and caloric models (Loesch et al. 2000; Vermillion & Lancaster 2022). The Collective is applying movement models to efficiently classify stopover locations, estimate length of stay, and determine bird use days to inform requests from managers with Migratory Bird Joint Ventures and U.S. State and Canadian Federal wildlife management agencies.

## 3. Education and Outreach

Despite increasing public awareness of declining bird populations, many audiences may not be aware of the broad array of habitats shorebirds need during their migrations. The term "shorebird" may limit conservation efforts in the Midcontinent, as many people do not connect them to grasslands or interior wetlands. Shorebirds are sometimes wrongly assumed to have similar habitat needs as other waterbirds like gulls, egrets, rails, or waterfowl. Environmental education can lead to indirect and direct conservation outcomes (Ardoin et al. 2020) and as an outreach tool, tracking data offer strong visual and hands-on learning opportunities to gain support for shorebird conservation efforts.

### School curriculum

Educators at Grays Harbor National Wildlife Refuge in Hoquiam, Washington, USA requested geographic coordinates and track lines to support their 3^rd^ and 4^th^ grade Shorebird Education Programs. The Collective provided data from 17 individuals of five species detected in or near Grays Harbor Estuary. Educators incorporated these data into four lessons, allowing students to map and explore shorebird movements within the Grays Harbor estuary and across the Pacific Flyway. The activity allowed the students to enhance skills in geography and ecology while learning how small actions in their communities can affect a shorebird’s entire migration. Refuge educators visited multiple classrooms every month, reaching a total of 870 students across 14 schools with this lesson over the 2022–23 school year.

### Community outreach

Nature Canada, a non-governmental conservation organization, requested tracking data for a comic following the movements of Bico, a fictional Hudsonian Godwit, as she travels from Canada to Chile. The comic’s goal is to raise awareness and generate excitement about shorebirds that visit James Bay, Canada, a globally important stopover area. The Collective provided two Hudsonian Godwit tracks and summary statistics (e.g., stopover sites and tenures, average flight speeds) for an infographic at the end of the comic. Nature Canada posted the comic online (<u>link</u>) and distributed paper copies in six First Nations communities around James Bay. The Collective provided similar information to the Minnesota Board of Water and Soil Resources, a state soil and water management agency, to communicate the role that wetland restoration and water management projects have in protecting shorebird populations in Minnesota. The Board shared this with their staff and conservation partners.

### Connecting communities with shorebird movements

The Western Hemisphere Shorebird Reserve Network, a science-based, partnership-driven, conservation initiative for protecting shorebirds through a network of key sites, wanted to show how shorebird movements connect “sister sites” within their reserve network. The Collective created an online storytelling series using ArcGIS StoryMaps to follow three tracked individuals as they migrated across the Americas (http://tiny.cc/ssccstorymap). Each StoryMap features a migratory bird flyway (Pacific, Midcontinent, Atlantic) and cultural theme (food, music, crafts), highlighting the people and places each shorebird might encounter along their migrations.

## Successes and future

The Shorebird Science and Conservation Collective is advancing shorebird conservation through collaborative data sharing and analysis in support of needs identified by practitioners. Such on-the-ground conservation is especially crucial given long-term and, now, accelerating shorebird population declines (Rosenberg et al. 2019; Smith et al. 2023). During the first two years we worked to address limitations to data access and analysis, and to conservation implementation (Table 1) in order to advance data-driven shorebird conservation.

We created a central resource of over 3,300 individual movement paths of shorebirds and provided staff to facilitate data access, and interpret and analyze shorebird science. New data become available as research projects develop and new contributors join. Staff share scientific expertise with practitioners, including specialized knowledge of shorebird ecology and statistical approaches to analyzing tracking data. Through these free scientific services offered to practitioners, and through the Collective’s extensive outreach throughout the Midcontinent, we have made it easier for practitioners to learn how shorebirds use their jurisdictions and to consider shorebirds when implementing their job duties. Our structure facilitates interaction, leading to co-production of analyses to inform land conservation, species management, and education and outreach.

The Collective is also providing a more complete understanding of shorebird ecology across the Americas, but some data limitations remain. For example, data contributed to the Collective are limited in some places or for some species and some datasets have high spatial uncertainty or low sampling frequency. Small sample sizes can limit generalizability and statistical inference (Hebblewhite & Haydon 2010). These factors are inherent to collated tracking datasets (Block et al. 2011), requiring careful evaluation of the suitability of available data to specific conservation questions, and communication of uncertainty (something the Collective emphasizes with partners). However, even small datasets can provide meaningful insights (Sequeira et al. 2019), and tracking data are often the only data available for shorebirds on private lands. Tracking datasets are the foundation of the Collective’s ability to implement bird conservation, but monitoring programs (e.g., International Shorebird Survey, https://www.manomet.org/project/international-shorebird-survey/; Migratory Shorebird Project, https://migratoryshorebirdproject.org/) and eBird participatory science data complement tracking data to inform conservation, especially where tracking data are limited (often for small-bodied species). Integrating across data types and facilitating communication with data registry efforts (Rutz 2022; Nightingale 2023) to create efficient and effective pipelines from data to action is a priority.

Due to privacy settings for some datasets, the Collective has not made a public mapping tool. Complementary tools are available that allow practitioners to explore tracking maps (e.g., Bird Migration Explorer, Smith et al. 2022). Data repositories (e.g. Movebank, Davidson et al. 2022; Motus, Taylor et al. 2017) allow those with time and motivation to find and download open access data, or to request permission to download private data, but see limitations (Figure 1, Scarpignato et al. 2023). However, practitioners do not often request raw data from us, or typically have staff capacity, time, motivation, or background to find and analyze data, even if all data were open access. The Collective adds value by analyzing and interpreting a complex hemispheric-scale dataset of shorebird movements in a way that is understandable, co-developed, and delivered in preferred formats. This lowers the activation energy for practitioners to consider and use shorebird science.

Most projects supported by the Collective are in response to direct requests, referrals, or targeted outreach. There is a need for future contributions to inform regional, national, and international priorities through strategic analyses designed to have the biggest effect on reversing shorebird population declines. We recognize potential to inform policy and funding decisions with shorebird science. For example, in the United States, federal agencies are administering landmark investments in climate and conservation programs through the 2022 Inflation Reduction Act (H.R.5376). The Collective’s consolidated dataset provides spatial and temporal evidence of shorebird habitat use that is also critical for informing international conservation conventions and agreements (Navedo & Piersma 2023).

Our funding prioritized the Midcontinent as a proof-of-concept, but since the Collective’s inception, partners have emphasized the importance of taking a full annual cycle approach to shorebird conservation. While our highest engagement came from practitioners in Texas, USA, the Collective also supported projects in the Pacific and Atlantic Flyways of North America, and data were contributed from scientists in 14 countries. We recognize an urgent need to secure additional funding to continue existing support of conservation, and to expand the Collective’s impact in the Caribbean region and Central and South America. Priorities include hiring specialists focused on supporting conservation initiatives in these regions, and who are fluent in Spanish, Portuguese, and French, and the inclusion of tracking data from species endemic to Latin America and the Caribbean. The Collective’s synthesized dataset can also help prioritize research to fill knowledge gaps. For example, tracking data are already being used to help inform shorebird surveys of the Amazon Basin, where eBird and survey data are extremely limited.

Innovations in tracking technology continue to enhance our understanding of shorebird migrations and habitat use—patterns that are changing with persistent habitat loss and climate change (Sutherland et al. 2012; Galbraith et al. 2014). Ongoing data collection and sharing are thus vital for liaisons like the Collective to rapidly integrate and interpret research for conservation in a changing world.

## Acknowledgments

Support for the Shorebird Science and Conservation Collective is provided by the Knobloch Family Foundation. Additional support is provided from ConocoPhillips Charitable Investments in support of the Migratory Connectivity Project (A.-L.H.) and from the U.S. Fish and Wildlife Service (R.B.L.). Thanks to the Advisory Group (Appendix I) and for data or conservation contributions from B. Ballard, B. Corey, G. Castresana, K. De Santiago, P. Erdmann, J. Pinnix, D. Ruthrauff, R. Summers, J. Shackelford, D. Shaw, M. Singer, T.L. Tibbitts, M. Weegman, L. Wright. Thanks to all who joined listening sessions and provided early feedback. Finally, thanks to funders, collaborators, volunteers, and staff who collected or supported the collection of shorebird tracking data. Any use of trade, product or firm names is for descriptive purposes only and does not imply endorsement by the U.S. Government.

## Notes

### Competing Interest Statement

The authors have declared no competing interest.

### Summary of Updates

Original pre-print prior to peer-review together with full appendices.

