## Supplemental Information for "The collective application of shorebird tracking data to conservation"

### **Appendix I:** Shorebird Collective Advisory Group

**Brad Andres,** U.S. Fish and Wildlife Service, National Shorebird Coordinator (Retired)

**Tjalle Boorsma,** Asociación Armonía

**Stephen Brown,** Manomet, Inc.

**Rob Clay,** Manomet, Inc., Western Hemisphere Reserve Network

**Ashley Dayer,** Virginia Polytechnic Institute and State University

**Jill Deppe,** National Audubon Society

**Jim Devries,** Ducks Unlimited Canada

**Eduardo Gallo-Cajiao,** University of Washington, Seattle

**Richard Gibbons,** American Bird Conservancy

**Catherine Hickey,** Point Blue Conservation Science

**Scott Johnston,** U.S. Fish and Wildlife Service, Northeast Region (R5)

**Matt Reiter,** Point Blue Conservation Science

**Stan Senner,** National Audubon Society (Retired)

**Paul Smith,** Environment and Climate Change Canada

**Kelly Srigley Werner,** U.S. Fish and Wildlife Service, Partners for Fish and Wildlife Program (Retired)

**EJ Williams,** American Bird Conservancy

### Appendix II: Data Sharing Agreement Form

#### Shorebird Science & Conservation Collective

*Data Contributor and License Agreement*

Link to online DSA webform: https://nationalzoo.si.edu/migratory-birds/shorebird-science-and-conservation-collective-data-sharing-agreement-form

Questions? Please contact Autumn-Lynn Harrison at

#### Background information and Detail of the Agreement

The Shorebird Science and Conservation Collective (hereafter, "Collective" or "Shorebird Collective") is a partnership of scientists and practitioners working to translate the collective findings of shorebird scientists into effective on-the-ground conservation to help reverse the decline of shorebirds across the Western Hemisphere. We pledge to do this with transparency regarding how data are used to support conservation.

Founded in late-2020 with three years of funding provided by the Knobloch Family Foundation, the work of the Collective is accomplished through three Knobloch Shorebird Conservation Fellows (hereafter Knobloch Fellows) based at the Smithsonian Migratory Bird Center that work with stakeholders across the Western Hemisphere. The Knobloch Fellows use available (i.e., from the literature or in the public domain) and contributed data to provide analytical and scientific support to multiple conservation and management stakeholders. An Advisory group composed of shorebird conservationists and researchers help guide the focus and priority projects of the Collective to ensure that the work of the Collective complements and supports existing efforts undertaken by individual scientists and organizations. A small amount of funding is also available to implement a limited number of projects on the ground.

For an introduction to the idea of the Collective and its planned work, a recorded webinar is available here: https://bit.ly/3601F1D

We invite you to join the Shorebird Collective as a Data Contributor and Science Partner. Our hope is that our plan for including your data will provide allowances for a range of data-sharing permissions depending upon contributor needs (see below).

##### Ways your data may be used depending upon the permissions you set:

**1. Data exploration:**

Prior to use of data in any conservation applications or outreach materials (see options below), the Collective will first explore the data to determine its suitability for specific applications. This would include work to determine the time frame of the data, the geographic location of the data, data quality, data gaps, preliminary analyses, etc. This may result in your data not being used. The Collective will, however, send all generated products to the Data Contributor (e.g., modeled datasets, processed geolocator datasets, etc.) upon request.

**2. Conservation applications and outreach products:**

We may compile, map, and or analyze the data you share to answer specific conservation questions. Maps, animations, and other visuals may be used in conservation workshops and/or for outreach products to illustrate conservation issues or concepts to stakeholders.

Three example conservation questions are below that were submitted by partners to the Collective:

- **Where should habitat conservation investments be made to benefit specific Canadian shorebird breeding populations?** Example products: Connectivity and site use maps that can be searched by species (and groups of species), sub-species, population, and geography and linked to population trends to help determine where conservation dollars should be spent.

- **The prairie potholes region is difficult to accurately survey for shorebirds. From tracking data could you tell us the places that are used and the times shorebirds visit so we can target our habitat conservation work and landowner training?** Example products: Reports and outreach products (PowerPoint, handouts) that can be delivered to managers during workshops and training programs; these products might include animations of tracking data (if contributors agree) as outreach tools to aid in landowner training; data on the timing and duration of use to support applications for sites to become part of the Western Hemisphere Reserve Network or other conservation programs.

- **I am considering placing some of my ranch holdings into a conservation easement. Can you use shorebird tracking and count data to identify the best places for me to set aside for shorebirds?** Example products: Planning exercises to identify lands in need of conservation that would most benefit shorebirds.

**3. Websites and social media channels for educational and outreach purposes:**
We may share conservation or outreach products on websites and social media channels for educational and outreach purposes, for example, on the Smithsonian Institution website, the Western Hemisphere Shorebird Group website, Shorebird Collective social media feeds, and other uses by request of Shorebird Collective partners.

**4. Reports to funders:**
The Collective will provide reports to funders that may include maps or images of outreach products.

**5. Scientific publications:**
Although not the primary focus of the Collective, we anticipate there may be opportunities for scientific publications, for example, methods papers, synthesis papers, or applied conservation papers.

#### You can decide how to share your data:

To participate in the Collective, a Data Contributor provides permission to the Collective to use the data for all exploratory uses (Use Category 1 above) without additional approval. The contributor will be provided with any products or analyses produced upon request.

For use category 2 above (Conservation Applications), you can specify one of two options:

a) The Collective may use the data for all uses without additional approval ("All uses are pre-approved"). The contributor will receive a notification of this use and will be provided with any products or analyses produced.
b) For each unique use, a Collective representative shall describe the intended use to the contributor and request specific permission for that use. The contributor will be provided with any products or analyses produced.

For use categories 3 and 4 above, you can specify one of three options:

a) The Collective may use the data for all uses without additional approval ("All uses are pre-approved"). The contributor will receive a notification of this use and will be provided with any products or analyses produced.
b) For each unique use, a Collective representative shall describe the intended use to the contributor and request specific permission for that use. The contributor will be provided with any products or analyses produced.
c) The data are not available for this category of use at this time. The contributor may specify an available date by which the data would be available (e.g., 3 years after data has been contributed).

For use category 5 above (scientific publications), you can specify one of two options. However, please note that data will never be used in a publication without permission, and we commit to early involvement of contributors in publication ideas. The two options are:

a) For any ideas for publications, the Data Contributor will be invited to participate early in the process. This may occur by email, or through a webinar with multiple potential authors. The Data Contributor can then decide whether they would like to participate as a co-author and have their data included.
b) The data are not available for this category of use at this time. The contributor may specify an available date by which the data would be available.

#### Data Ownership

Data contributed to the Shorebird Science and Conservation Collective remains the property of the data owner at all times and will not be shared with any third party without the previous agreement of the Data Contributor as noted below. If the Data

Contributor would like to discontinue working with the Collective in the future, they may notify the Collective in writing (by email or letter) to revoke any permissions previously granted. This will nullify any previously signed Data Sharing Agreements.

#### Who will work with your data and how will it be protected?

Data sharing agreements with Data Contributors will be managed by staff supporting the Collective at the Smithsonian Migratory Bird Center. To transfer tracking data, contributors will be encouraged to share their data via Movebank (www.movebank.org) and to add the Shorebird Collective as a collaborator under sharing permissions. If Data Contributors do not wish to store their data in Movebank, they may also transfer the raw tracking data and associated metadata to the staff supporting the Collective at the Smithsonian Migratory Bird Center. Staff who may work with your data for the purposes of the Collective include:

- Autumn-Lynn Harrison –Research Ecologist with the Smithsonian Migratory Bird Center
- Three Knobloch Shorebird Conservation Fellows

Data may also be shared with contractors, interns, students, or Collective collaborators solely for purposes of working on products designed for the Collective, all under the direction of the Smithsonian staff supporting the Collective and under the stipulations of this data sharing agreement. No contributed data will be shared outside of the stated uses of the Collective unless the Data Contributor expressly grants permission for the contributed data to be shared with other initiatives. For example, data will only be shared with the National Audubon Society's Migratory Bird Initiative and the Smithsonian Institute's Migratory Connectivity Project (to produce the "Atlas of Migratory Connectivity for the Birds of North America") if copies of written agreements between the Data Contributor and these two other initiatives are provided.

#### For how long am I providing permission for data use?

The Collective is requesting permission to use the data indefinitely for the purposes describe above. No further use of the data will be permitted without your permission. Data contributors may choose to exit this agreement at any time and must notify a staff member of the Shorebird Collective by email.

#### How will my contribution be acknowledged?

Data Contributors will be acknowledged in all products that incorporate their data unless a contributor notifies the Collective that they wish not to be acknowledged. Data Contributors will be responsible for providing appropriate names, organizations, social media handles or other relevant citation materials for each dataset as needed. The Collective will maintain this information in our Data sharing permissions database.

In addition, the Collective will establish a "wall of conservation sharing" on its website that lists each project that has agreed to provide their data for the conservation of shorebirds. To help you measure the conservation impact of your scientific contributions, this website will also include a list of conservation projects that used the data. We think giving credit to Data Contributors for advancing shorebird conservation is critically important.

#### What are my responsibilities as a Data Contributor and Science Partner?

Data Contributors should act in good faith as a science partner to assist the Collective's staff in providing, understanding, analyzing, and applying their data so there is no misinterpretation of their data for conservation applications. Data Contributors shall inform the Collective of any issues with the timely submission of data, as well as updates on errors, inaccuracies or other problems with respect to their contributed data as soon as possible after becoming aware of such issues. Data Contributors are also obligated to provide regular updates on the best way for the Shorebird Collective to communicate with them (including updates to e-mail addresses or other preferred means of communication). The Data Contributor must notify the Collective if ownership of the data transfers hands so communication with the new leader can be established.

Data Contributors are expected to respond in a timely manner to all requests for data uses that are not pre-approved according to the level of sharing agreed to in the submitted agreement. If no requested uses are approved by the Data Contributor in the first year of the Collective's work, the Data Contributor will be notified that they will no longer be considered a scientific partner in the Shorebird Collective, and we will no longer investigate uses of their data. Their names will also be removed from the "wall of conservation sharing" (see above).

In case of loss of contact with a data-owner (i.e., non-reply to e-mails from the Collective for more than 12 months), the Collective will manage the data request procedure on behalf of the data owner, consistent with any limitations originally stipulated by the Data Contributor.

##### Will I be notified of any revisions to this data sharing agreement?

As the Collective is a new initiative, we anticipate there may be a need to revise this agreement. For changes that do not affect the data use (i.e., to clarify potential uses), the Data Contributor will be notified without further action needed. All signatories will be notified of any changes that affect data use and can choose to opt out of future uses upon this notification.

##### May I use my own organization's data sharing agreement?

Yes. If your organization has a standard data use agreement, you may request that we sign it, provided it will allow use of the data for the purposes described above in the categories you select in the Collective's Data Sharing Agreement. Additionally, if you select data use options in any category that require users to seek permission before use, you are free to stipulate your own conditions in order to approve a specific data request.

#### Data Sharing Agreement & License for Use

A single form should be submitted for a given project (may include multiple species) by a designated lead contributor. Other individuals involved in this project should be listed below as "contributors of the dataset". If you have multiple types of data that you wish to share differently, please fill out a separate form in each case. However, we encourage you to share as much data as possible and to keep the agreements as simple as possible.

Last name, First name: *
Your answer

Job Title, Organization: *
Your answer

E-mail address/Phone number: *
Your answer

State or Province, Country of Organization:
Your answer

What is your preferred form of communication?
☐ Email
☐ Phone

Please provide a brief description of the dataset you are contributing (include information on species, location tags deployed, and question addressed). If your preferred data sharing permissions differ between species, then separate forms should be submitted for each species. *
Your answer

What tracking technology was used to collect the data? (list all types separated by species if different; if your preferred data sharing permissions differ between technologies then separate forms should be submitted for each technology) *
Your answer

##### For the following data uses (see above), please select the level of sharing you would like us to apply:

Data Exploration *

**What do we mean by data exploration?** During data exploration, we generate maps, figures, and tables that summarize the number of tracked birds and the broad distribution of data contributed to the Shorebird Collective in a region (e.g., state/province, county, ecoregion). The maps do not show raw tracks or points and are summarized across datasets to maintain data privacy. We show these products to conservation practitioners to determine if more detailed examination of data to support a conservation project might be worthwhile. These maps, figures, and tables may also be incorporated into conservation reports, presentations, newsletters, and our website. You will always be acknowledged for this data contribution. For any data uses outside of the summary maps, figures, and tables, we follow terms specified in data use agreements and will reach out directly to obtain permission if required in the DSA.

☐ By contributing data to the Shorebird Collective, I agree to allow my data to be explored for conservation applications as described above.

Conservation Applications and Outreach Products *
☐ I pre-approve all uses
☐ Please notify me prior to each use

Web and Social media Sharing for sharing news of the Collective and for educational outreach *
☐ I pre-approve all uses
☐ Please notify me prior to each use
☐ Do not use for this purpose at this time

Funder Reports *
☐ I pre-approve all uses
☐ Please notify me prior to each use
☐ Do not use for this purpose at this time

Scientific publications *
☐ Please notify me early in the process when considering how my data might contribute to an idea for a scientific publication and to request permission for use.
☐ Do not use for this purpose at this time

Connecting contributors with researchers
The Shorebird Collective has received requests from scientists (including graduate students) seeking relevant data for research projects. The research may have a conservation motivation fitting the priorities of the Shorebird Collective's mission, but the requests also have a primary goal of publishing a peer-reviewed paper or contributing to a dissertation or thesis that would require additional involvement and permissions from data contributors to the Collective. The Shorebird Collective is currently prioritizing supporting requests that come directly from conservation and management communities working on-the-ground. We have been guided by our Advisory Group to act as a connector/data discovery service for research requests, rather than to facilitate permissions and data sharing. For these types of requests, please specify if you allow the Shorebird Collective to share your name and contact information with researchers. If you approve of sharing your contact information with the requesting scientist, the scientist will then reach out directly to you to propose the potential research collaboration.

Please select from one of the following:
☐ I agree that the Shorebird Collective may share my name and email address with requesting researchers.
☐ Please do not share my name or email with requesting researchers.

Please list any additional contributors (i.e., co-owners) of the dataset.
Your answer

I have checked with other contributors and all have given permission for data to be contributed and used as indicated in this agreement: *
☐ Yes
☐ Not yet

How will you be sharing your data? If other, please describe below. *
☐ With Movebank (Our Movebank username is ShorebirdCollective)
☐ Direct Send
☐ Other:

If you will share your data with Movebank, please provide your username:
Your answer

If these data have been published, please provide full citation.
Your answer

If your data have not yet been published, how would you like the data to be cited? (e.g., Contributor name, Project name, and Organization)?
Your answer

#### Additional conditions

Please describe any additional conditions that must be met for this data sharing agreement to be finalized. If the condition is outside the scope of this agreement, an amendment may be necessary.
Your answer

#### Agreement

By submitting this form, you grant a non-exclusive, royalty-free license to the Smithsonian Migratory Bird Center to use, in whole or part, your contributed data in connection with the Shorebird Science and Conservation Collective as indicated in the data use section above. By contributing data, Data Contributors and the Shorebird Collective team agree to adhere to the above terms of use.

Please contact Autumn-Lynn Harrison with questions.

#### Please click to indicate your agreement *

☐ I have read and agree to the terms of the Shorebird Science and Conservation Collective data contributor and license agreement

### **Appendix III:** Conservation Use Request Form for Practitioners

#### Shorebird Science & Conservation Collective

*Request for Use of Shorebird Tracking Data*

Smithsonian
*Migratory Bird Center*

Link to online data request webform: https://nationalzoo.si.edu/migratory-birds/request-for-use-shorebird-tracking-data

Collaborators across the Western Hemisphere have contributed tracking data from over 2,800 individuals representing 31 shorebird species as a single resource for use in management, conservation, and education and outreach. Find out more here: https://nationalzoo.si.edu/migratory-birds/shorebird-collective.

If you have a project that could benefit from this resource, please use this form to provide details of your request. Requests should be specific and provide enough context for Shorebird Collective team members to understand the request. Completed forms will be reviewed by the Shorebird Collective and a team member will reply with additional questions and/or updates as needed. If you have any questions, please contact Regional Conservation Specialist, Candace Stenzel,.

***What data can the Shorebird Collective provide?***
The Shorebird Collective team based at Smithsonian's National Zoo and Conservation Biology Institute are the curators of the contributed data, but the data remained "owned" by the contributors and use is governed by a Data Sharing Agreement. We facilitate data discovery and requests for use. In many cases, contributors have provided permission for use in conservation or outreach without needing further requests. In other cases, we may require additional approval, especially for requests for raw data or use in scholarly work (for example, in scientific publications). Requests are therefore not guaranteed.

The Shorebird Collective can provide maps and summary statistics for shorebird tracking data relevant to your project. A Shorebird Collective team member may also be able to collaborate with you on a needed analysis for your conservation project depending on time and priorities.

If you would like to learn more about the Shorebird Collective and what data and resources may be relevant to your work before completing this form, please contact Regional Conservation Specialist, Candace Stenzel,.

First and Last Name:
*Answer*

Job Title, Organization:
*Answer*

State/Province, Country of Organization:
*Answer*

E-mail address:
*Answer*

Phone number:
*Answer*

Names and Organizations of any co-requesters to this data request:
*Answer*

Please provide a brief overview of your request, with a general description of what information is needed, for what species and areas, and for what purposes.
*Answer*

Please provide details of who will use and/or view any data or products provided by the Shorebird Collective and its contributors.
*Answer*

What end-product are you requesting (e.g., map, summary statistics, written description, other)?
*Answer*

How will the end-product be distributed? How many people will the end-product reach?
*Answer*

What is your timeline for receiving products for your request?
*Answer*

### **Appendix IV:** Data Use Request From to Contributors

#### Shorebird Science & Conservation Collective

*Data Use Request for Conservation Application*

In your Data Use Agreement with the Shorebird Collective (**DSA_ID** submitted by **Name**, **Organization** on **MM/DD/YYYY**, focused on **SPECIES**) you indicated that you would like to be contacted for permission to use your contributed data for conservation application purposes. We now have a relevant request that we seek your permission for use. Details of the request are provided below. If you have any questions, please contact Regional Conservation Specialist, Candace Stenzel,.

| | |
|---|---|
| **Name of Project:** | |
| **Conservation Partners:** | |
| **Requested Data:** | *See attached map of **Species** tracks. |
| **Project Summary**: | |
| **Intended Outputs:** | |
| **Citation:** | Tracks will be attributed according to the citation you provided in your data sharing agreement:<br>• Data contributed by<br>• Data Co-owners:<br>• Data Owner Organizations:<br>• Citation: |

__Data Use Agreement__

☐ I hereby grant the Shorebird Collective to use these data for the purposes as described above.

☐ I do not wish to allow these data to be used for this purpose at this time.

**If applicable, please describe any additional conditions that must be met for the purposes of using the data as described above.**

Printed Name of Data Contributor

Signature of Data Contributor

Date

### **Appendix V:** Notification of Data Use Form to Contributors

#### Shorebird Science & Conservation Collective

*Notification of Data Use: Conservation Application*

Smithsonian
*Migratory Bird Center*

In your Data Use Agreement with the Shorebird Collective (**DSA_ID** submitted by **Name, Organization** on **MM/DD/YYYY**, focused on **SPECIES**), you provided preapproval to use the contributed data for conservation application purposes. This is a notification to inform you that we now have a relevant request for use of these data. Details of the request are provided below. If you have any questions, please contact Regional Conservation Specialist, Candace Stenzel,.

| | |
|---|---|
| **Name of Project:** | |
| **Conservation Partners:** | |
| **Requested Data:** | *See attached map of **Species** tracks. |
| **Project Summary**: | |
| **Intended Outputs:** | |
| **Citation:** | Tracks will be attributed according to the citation you provided in your data sharing agreement:<br>• Data contributed by<br>• Data Co-owners:<br>• Data Owner Organizations:<br>• Citation: |

### **Appendix VI:** Notification of Data Use (Filled example)

#### Shorebird Science & Conservation Collective

Smithsonian Migratory Bird Center

*Notification of Data Use: Conservation Application*

In your Data Use Agreement with the Shorebird Collective (DSA 40 submitted by Autumn-Lynn Harrison, Smithsonian Migratory Bird Center on 11/02/2021, focused on Long-billed Curlews), you provided preapproval to use the contributed data for Conservation Application purposes. This is a notification to inform you that we now have a relevant request for use of these data. Details of the request are provided below. If you have any questions, please contact Regional Conservation Specialist, Candace Stenzel,.

| | |
|---|---|
| **Name of Project:** | Nueces Delta Fire Management |
| **Conservation Partners:** | Coastal Bend Bays & Estuaries Program (CBBEP) |
| **Requested Data:** | Tracking data for 3 Long-billed Curlews (LBCU).<br>Movebank ID for LBCUs: 46195209<br>Individual IDs: 141775, 154072, 154074<br><br>*See attached map of LBCU tracks. |
| **Project Summary:** | The CBBEP requests shorebird location data within Nueces Delta Preserve (NDP) and Mission River property (MR) in Nueces County, Texas to assess shorebird use of habitat in response to prescribed burns and water level management. These data will support a new initiative focused on refining management techniques for shorebirds. This request is purely for internal summarization and consideration and will not be distributed outside of CBBEP. The birds listed above were detected in the NDP and/or MR property. The Shorebird Collective will be sharing location data of these birds with the CBBEP to support their new initiative. |
| **Intended Outputs:** | The following data will be shared with the CBBEP:<br>• Points and tracks for the 3 curlews detected in the NDP and MR property.<br>• Points and tracks for these birds in Texas to determine where and when they were moving to the NDP and MR (to compare to the timing of the controlled burns).<br>• Full annual track lines (points not needed) to show connections of the birds to other areas. |
| **Citation:** | The track will be attributed according to the citation you provided in your data sharing agreement:<br>• Data Contributor: Autumn-Lynn Harrison[1]<br>• Data Co-owners: David Newstead[2]<br>• Unpublished data, Migratory Connectivity Project<br>• Contributor Organizations: [1] Smithsonian Migratory Bird Center, [2] Coastal Bend Bays and Estuaries Coastal Bird Program[3] |

### Long-billed Curlew Tracks

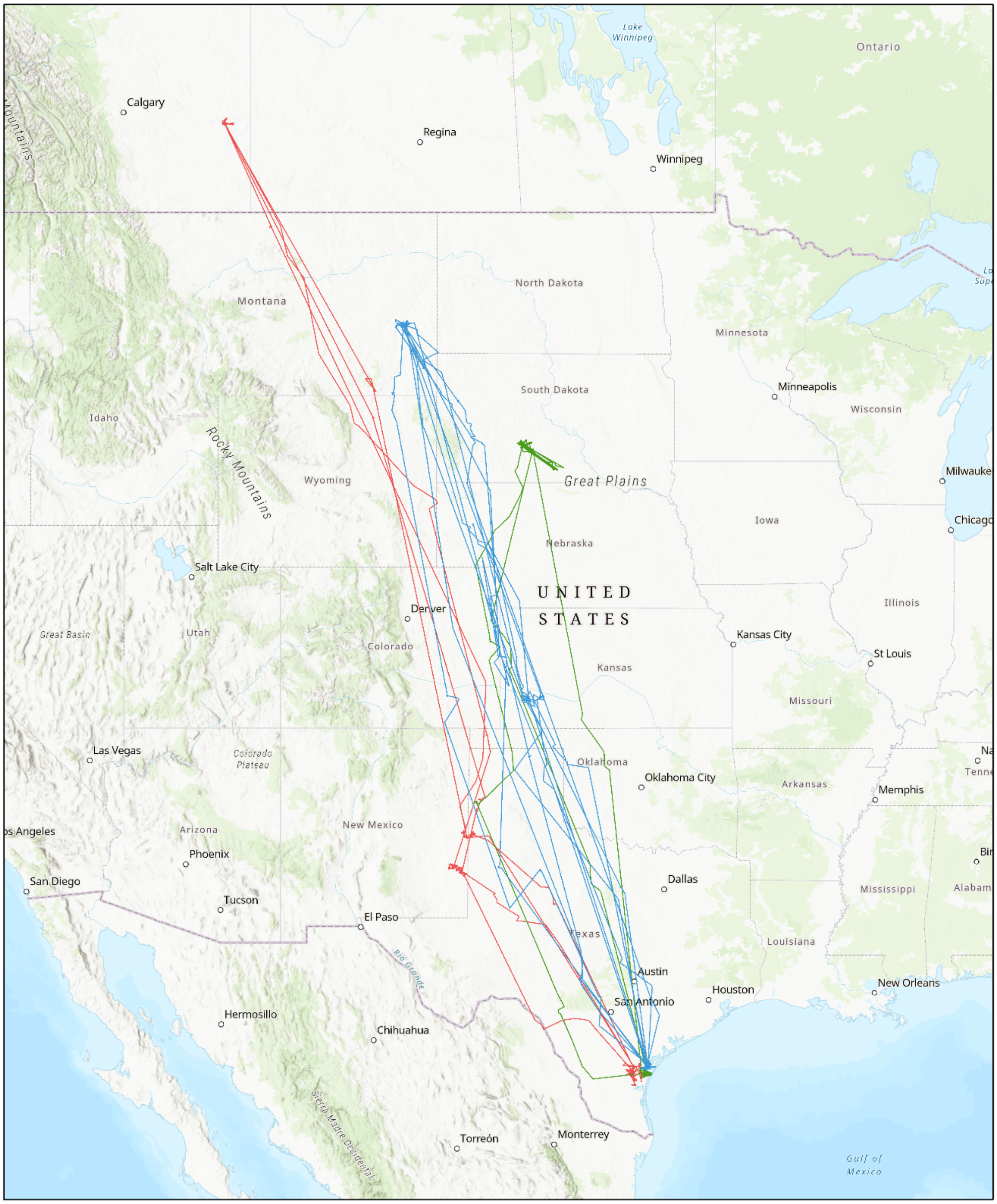

Individual Bird ID

- 141775
- 154072
- 154074

1:12,709,124

0 112.5 225 450 mi

0 180 360 720 km

Esri, HERE, Garmin, FAO, NOAA, USGS, EPA, Esri, USGS

### **Appendix VII:** Species List

Current list of shorebird species contributed to the Collective's dataset.

| **Common Name** | **Scientific Name** |
|---|---|
| American Golden Plover | *Pluvialis dominica* |
| American Oystercatcher | *Haematopus palliatus* |
| American Woodcock | *Scolopax minor* |
| Bar-tailed Godwit | *Limosa lapponica* |
| Black Oystercatcher | *Haematopus bachmani* |
| Black Turnstone | *Arenaria melanocephala* |
| Black-bellied Plover | *Pluvialis squatarola* |
| Bristle-thighed Curlew | *Numenius tahitiensis* |
| Buff-breasted Sandpiper | *Tryngites subruficollis* |
| Common Ringed Plover | *Charadrius hiaticula* |
| Dunlin | *Calidris alpina* |
| Greater Yellowlegs | *Tringa melanoleuca* |
| Hudsonian Godwit | *Limosa haemastica* |
| Least Sandpiper | *Calidris minutilla* |
| Lesser Yellowlegs | *Tringa flavipes* |
| Long-billed Curlew | *Numenius americanus* |
| Long-billed Dowitcher | *Limnodromus scolopaceus* |
| Marbled Godwit | *Limosa fedoa* |
| Mountain Plover | *Charadrius montanus* |
| Pacific Golden Plover | *Pluvialis fulva* |
| Pectoral Sandpiper | *Calidris melanotos* |
| Piping Plover | *Charadrius melodus* |
| Purple Sandpiper | *Calidris maritima* |
| Red Knot | *Calidris canutus* |
| Red Phalarope | *Phalaropus fulicaria* |
| Red-necked Phalarope | *Phalaropus lobatus* |
| Ruddy Turnstone | *Arenaria interpres* |
| Sanderling | *Calidris alba* |
| Semipalmated Plover | *Charadrius semipalmatus* |
| Semipalmated Sandpiper | *Calidris pusilla* |
| Short-billed Dowitcher | *Limnodromus griseus* |
| Solitary Sandpiper | *Tringa solitaria* |
| Upland Sandpiper | *Bartramia longicauda* |
| Western Sandpiper | *Calidris mauri* |
| Whimbrel | *Numenius phaeopus* |
| White-rumped Sandpiper | *Calidris fuscicollis* |
| Willet | *Tringa semipalmata* |
| Wilson's Plover | *Charadrius wilsonia* |
